# Molecular Dynamics simulation of TDP-43 RRM in the presence and absence of RNA

**DOI:** 10.1101/2022.03.15.484514

**Authors:** David Donald Scott, David Mowrey, Karthi Nagarajan, Liberty François-Moutal, Anil Nair, May Khanna

**Author notes:** To whom correspondence should be addressed: Dr. May Khanna, Department of Pharmacology, College of Medicine, University of Arizona, 1501 North Campbell Drive, P.O. Box 245050, Tucson, AZ 85724, USA; Dr. Anil Nair; Department of in silico Drug Discovery, Icagen – a Ligand Company, Ligand Pharmaceuticals, 12470 N Rancho Vistoso Blvd., Ste #150, Oro Valley, Tucson, AZ 85755. Authors contributed equally (David Donald Scott, David Mowrey, Karthi Nagarajan).

## Abstract

Structural characterization of the prion prone TAR DNA Binding protein (TDP)-43 has been challenging since its intrinsically disordered regions represents 15-30% of the total protein. TDP-43 is a nucleic acid binding protein with an N-terminal domain, two RNA Recognition Motifs (RRM1 and RRM2) and the C-terminal domain. In this study, we seek to define possible new targetable sites on the apo structure of TDP-43 RRM domains. To do so, we used molecular dynamic (MD) simulations on the NMR solved TDP-43_RRM1-2_ structure bound to RNA to predict the apo structure. Contact analysis of TDP-43 showed that while the integrity of the individual domains was maintained upon RNA removal, a decrease in interdomain contacts was observed. Moreover, we compared apo TDP-43 structures obtained from MD to AlphaFold 2 (AF2) predicted TDP-43 structures and found differences in loop regions. A Sitemap analysis identified five druggable sites for the RNA bound structure solved by NMR, while fewer sites were identified following MD simulations and AF2 predicted apo structures.

## Introduction

TAR DNA-binding protein 43 (TDP-43) is a RNA-binding protein that is nuclear, is implicated in RNA metabolism reported to be present as cytoplasmic inclusions in the neurons and/or glial cells of a range of neurodegenerative diseases including ALS-FTLD and AD^*1*^. Depletion of cellular TDP-43, as a result of being trapped in inclusions, leads to the loss of neuronal activities regulated by TDP-43 which further results in neuronal death^*2*^. TDP-43 exhibit multiple functional domains, can bind more than 6000 RNA transcripts^*3*^, and may adopt higher order of organization such as dimers, necessary for its splicing function^*4*^. TDP-43 can also form higher order reversible condensates, termed LLPS (for liquid-liquid phase separation)^*5*^. TDP-43 subdomains consist of an N-terminal domain (NTD), two RNA Recognition Motifs (RRM1 and RRM2) and the C-terminal domain (CTD). While the CTD is mostly unstructured, the NTD and RRM modules are well structured domains^*4, 6*^.

Due to TDP-43-centric pathologies in neurodegenerative diseases (NDs), this protein is a target of choice for therapeutic intervention. However, the localization – NTD, RRM or CTD – and the precise “form” or “state” of TDP-43 – RNA bound, RNA free, monomer, condensates, or aggregates – to be targeted to modulate TDP-43 cytotoxicity remain to be determined. We have developed small molecules targeting TDP-43 NTD as well as the RNA-bound RRM domains^*7, 8*^; interestingly both molecules reduce RNA binding to various extent and mitigate ALS phenotypes in Drosophila, although further characterization of the compounds are needed. Defining small molecules that can distinguish and target RNA-bound vs RNA-free forms of TDP-43 of the RRM domain, would help in evaluating through chemical biology which form of TDP-43 can be targeted and more importantly which form reduces neurotoxicity. However, to date, there is no apo structure of the RRM domains in tandem solved either by NMR or X-ray, likely due to the presence of a highly flexible loop (178-191), that confers adaptability to different nucleic acid partners by allowing different orientations of the RRM domains^*9*^. Molecular dynamics (MD) offers the possibility to study dynamics and to integrate particle motions, more particularly long-range inter-domain correlation motions. A recent study on the tethering of TDP-43 RRM domains used MD simulations, among other techniques, to show higher conformational dynamics as well as higher destabilization of TDP-43_RRM1-2_ compared to the individual domains^*10*^, further highlighting the dynamic nature of TDP-43.

In this study, we used MD simulations on the NMR solved TDP-43_RRM1-2_ structure (PDB ID: 4bs2^*6*^) to predict the apo TDP-43 or RNA-free structure. Contact analysis of TDP-43 showed that while the integrity of the individual domains was maintained upon RNA removal, a decrease in interdomain contacts was observed. We then compared apo TDP-43 structures obtained from MD simulations to AlphaFold2^*11*^ (AF2) predicted structures and found differences in loop regions. Finally, a Sitemap analysis showed five druggable sites on the average of the RNA bound structure solved by NMR, while only three pockets were found for both MD and AF2 predicted apo structures.

## Experimental Procedures

### Molecular Dynamics of Apo-TDP43

Five structures were selected from the NMR ensemble of TDP43 (PDB ID: 4bs2^*6*^) based on energy and structural diversity. For each structure, we performed three 1 μs molecular dynamics simulations using three different random seeds. Simulations were performed both with and without the RNA present in the simulation using Desmond with the OPLS3e forcefield^*12, 13*^. Each simulation system was immersed in a cubic water box with a 1 nm buffer distance. The system was neutralized and included 150 mM NaCl. Prior to production runs, each system underwent 100 ps of Brownian dynamics simulation (NVT, 10 K). Restraints were gradually released over 24 ps of NPT simulation (1 bar, 310 K). We used 9 Å cutoffs for VDW and coulombic interactions and a 2 fs timestep. Simulation frames were saved every 0.1 ns. All analyses, unless otherwise noted, were performed on the combined frames of all 15 simulations (5 structures × 3 random seeds) sampled every 0.2 ns. For root mean square fluctuation (RMSF) and cross-correlation analysis, reference residues used for the alignment were those contained within secondary structural elements minus those with an RMSF greater than three standard deviations from the mean.

For contact map and cross-correlation analysis the first 100 ns of simulation (500 frames) of each simulation were discarded and all 15 simulations were concatenated. A contact between residues *i* and *j* was assigned if any heavy atom of residue *i* is within 4.5 Å of any heavy atom of residue *j*. Contacts were only considered in the final analysis if the interaction was present for >= 39% of the concatenated trajectory (p<0.01 from a one-sided test from zero). Post-hoc analysis was performed computing the p-value on the difference between the means of the two Bernoulli distributions.

For interdomain angle analysis the first 100 ns of simulation (500 frames) were also discarded and the remaining frames of all 15 simulations were concatenated. To measure relative orientations between the two domains we measured both a dihedral twisting angle (τ) and an RRM1 opening angle (α). For the τ angle we chose residues G148 and A230 as the center of each domain and residues C173 and R191 to define the axis of rotation. The full dihedral angle is defined using Cα atoms of the residue list G148 – C173 – R191 – A230. The α angle was measured using Cα atoms of the residue list G148 - C173 - R191 (Fig. 1a).

**Fig. 1.**
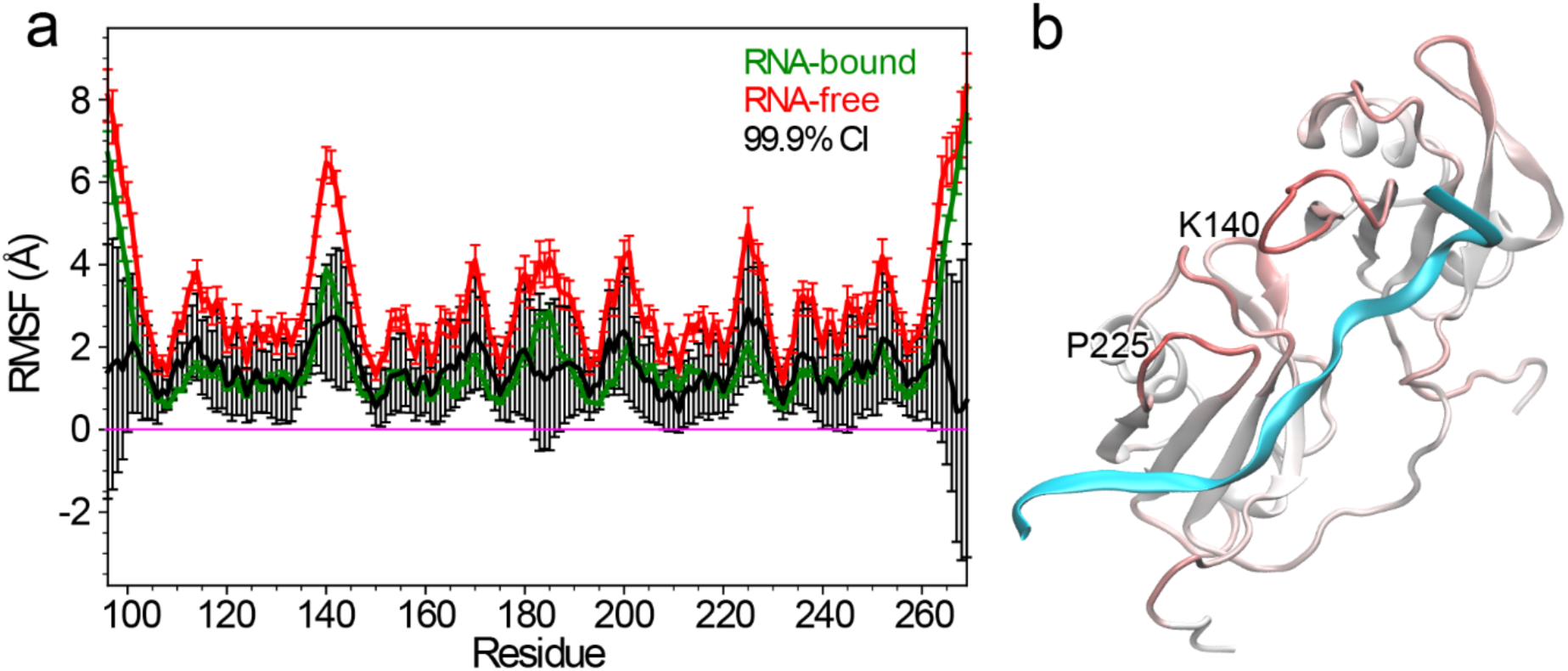
Differences in root mean square fluctuation (RMSF) between RNA-free and RNA-bound simulations of TDP43. (**a**) Plot of the RMSF values for RNA-free (red) and RNA-bound (green) simulations. Error bars are the standard error among 15 simulations. The black lines represent the 99.9% confidence interval for the difference between means. (**b**) The structure of TDP43 colored by difference in RMSF between RNA-free and RNA-bound from white (minimal RMSF difference) to red (maximum RMSF difference). The RNA strand is shown as a cyan ribbon for reference. Nearly all residues of RNA-free TDP43 show a significant increase in fluctuations compared to the RNA-bound TDP43, exceptions being the N- and C-termini and a short span between the two domains.

### AlphaFold 2 prediction of apo TDP-43 RRM domains

Computational approaches -first with AlphaFold then with AlphaFold 2 (AF2)^*14*^, a neural network-based algorithm that integrates multi-sequences alignments and training procedures based on the evolutionary, physical and geometric constraints of protein structures- are now able to predict protein structures with high accuracy. We thus used AF2 to predict apo TDP-43 RRM domains (**Fig. S2**) and obtained five models (named here AF_1-5). Only the RRM domains (102-269 amino acids) construct was submitted to structure prediction.

### Sitemap Analysis

Sitemap analysis was done using Schrodinger^*15*^ as previously described^*8*^. We attempted to converge an average structure from the multiple structures generated by MD simulations and AF2 models, but they resulted in non-physical structures. Therefore, the best representative structure based on minimal energy refinement was used.

## Results

To assess the system equilibration the root mean square deviation (RMSD) between the starting structure of the simulation and all succeeding frames was assessed (**Fig. S1**). Plots of the RMSD over time show that in most cases (13 out of 15 simulations) the trajectories of RNA-bound simulations equilibrate within 50 ns of simulation while the other two simulations show a slight jump later in the trajectory^*10*^. In contrast, simulations of the RNA-free system showed much greater deviation from the starting structure and required significantly more time to reach equilibrium in this case reaching a steady trajectory after 600 ns of simulation.

### Root-mean-square-fluctuation (RMSF) Analysis

Root-mean-square-fluctuation (RMSF) analysis measures the positional variance of each residue over the course of the trajectory and indicates residue flexibility over the course of the trajectory (**Fig. 1a**). As expected and consistent with the RMSD analysis, we observe a significant global increase in backbone flexibility in the absence of bound RNA. Particularly large increases in flexibility occur around residues K140 and P225, located in loop regions (**Fig. 1b**).

We next performed contact analysis of TDP-43 to assess any global changes or rearrangements in structure upon removal of RNA from the initial NMR structure (**Fig. 2**). The contact maps show that the structural integrity of individual domains is maintained in the absence of bound RNA (**Fig. 2a and b**). However, we observe a greater decrease in interdomain contacts as compared to formation of new contacts. Specifically, we observe significant decreases in contact probability (p<0.05) between residue pairs D247-S104, D247-D105, R151-D247, I249-L131, I249-M132, I249-R151, G252-M132, S254-D105, and S254-M132. Two new contacts are predicted to form between residues E261-K192 and F231-P223 (**Fig. 2c**).

**Fig. 2.**
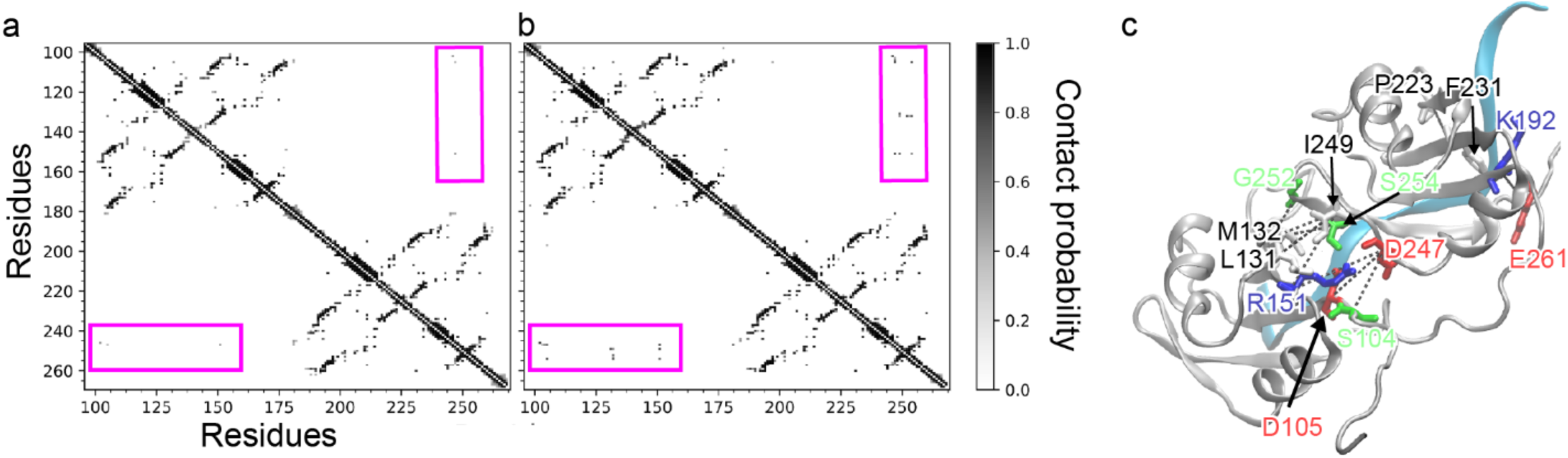
Contact maps of RNA-free (a) and RNA-bound (b) TDP43. Contact maps were computed from concatenating all 15 trajectories with frames sampled every 0.2 ns over the 1000 ns simulation (75000 frames total). A contact exists between residues *i* and *j* if any heavy atom of residue *i* is within 4.5 Å of any heavy atom of residue *j*. Contacts are only shown if the interaction is present for >= 39% of the concatenated trajectory (p<0.01 from a one-sided test from zero). Contact maps show that the structural integrity of each individual domain (upper left and lower right quadrants) is maintained between RNA-free and RNA-bound simulations. A major change between the contact maps is decreased interdomain contacts in the RNA-free simulations compared to RNA-bound simulations (lower left and upper right quadrants). (**c**) Residues showing significant changes in contacts between the two systems are highlighted as sticks on the TDP43 structure and colored by residue type white – nonpolar, green – polar, red – acidic, and blue – basic. The bound RNA is shown as a cyan ribbon. In the absence of RNA, the two domains are held less rigidly together and are freer to adopt other conformations with respect to each other.

### Intra- and Inter-domain Motion Analysis

Cross-correlation matrices were generated to determine changes in correlated motion between RNA-free and RNA-bound simulations (**Fig. 3**). Motion can be correlated (residues move in the same direction), anti-correlated (residues move in opposite directions), or uncorrelated (the movement of one residue is independent of the other residue). Cross-correlation matrices of RNA-free and RNA-bound simulations show much stronger correlations in RNA-free simulations than in RNA-bound simulations. Most of the increased correlated motion is likely the result of the overall increased motion in RNA-free simulations compared to RNA-bound simulations. In several instances, however, we observe increased interdomain correlated motion that significantly exceeds the increased intradomain correlated motion. The magnitude of the increase in intradomain correlated motion is likely indicative of the magnitude of the effect coming from an overall increase in motion. The fact that the interdomain correlated motion exceeds the intradomain correlated motion highlights its significance.

**Fig. 3.**
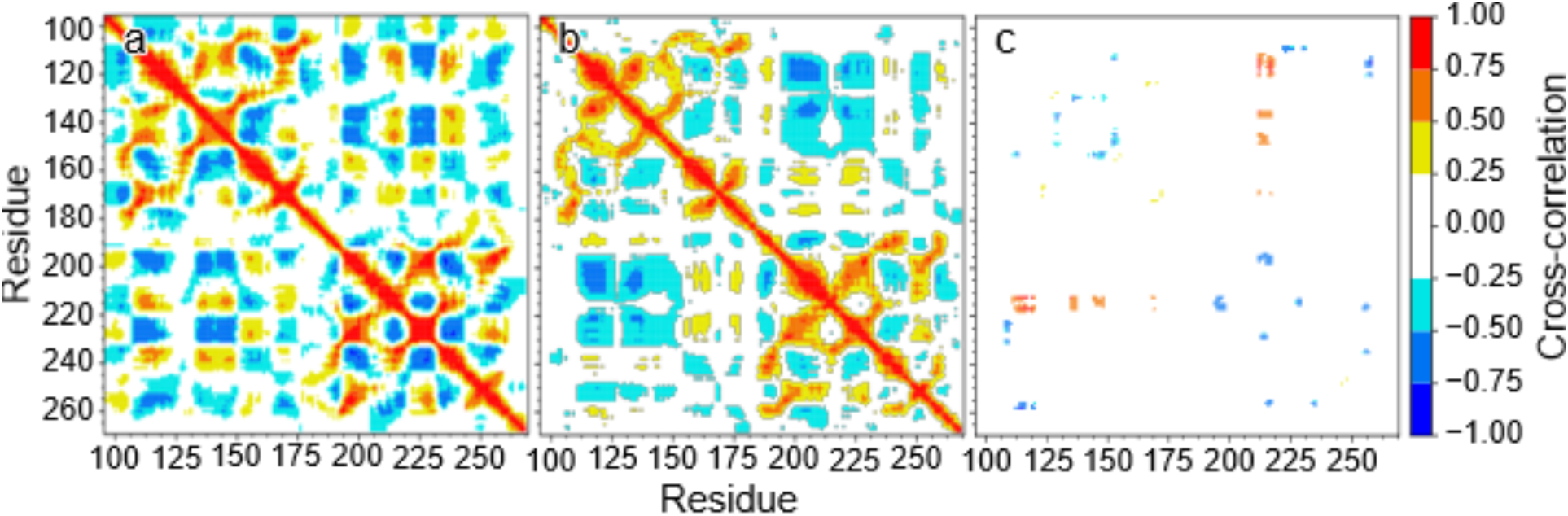
Cross-correlation matrices for RNA-free (a), RNA-bound (b), and the difference between RNA-free and RNA-bound (c). Cross-correlation matrices were computed from concatenating all 15 trajectories with frames sampled every 0.2 ns over the 1000 ns simulation (75000 frames total) based on Cα atoms. Significance filters were applied so only correlations passing a 5σ threshold are shown.

To further characterize inter-domain motion, we performed angle analysis examining the relative orientation of the two domains with respect to each other (**Fig. 4**). We measured both a dihedral twisting angle (τ) and an RRM1 opening angle (α). The analysis shows that in the absence of RNA there is a significant increase in orientational variability from what is observed in RNA-bound simulations. The increases come from variations in both angles; however, several distinct populations can be observed indicating the presence of multiple stable conformations. These results are consistent with increased inter-domain correlated motion in the absence of RNA-binding.

**Fig 4.**
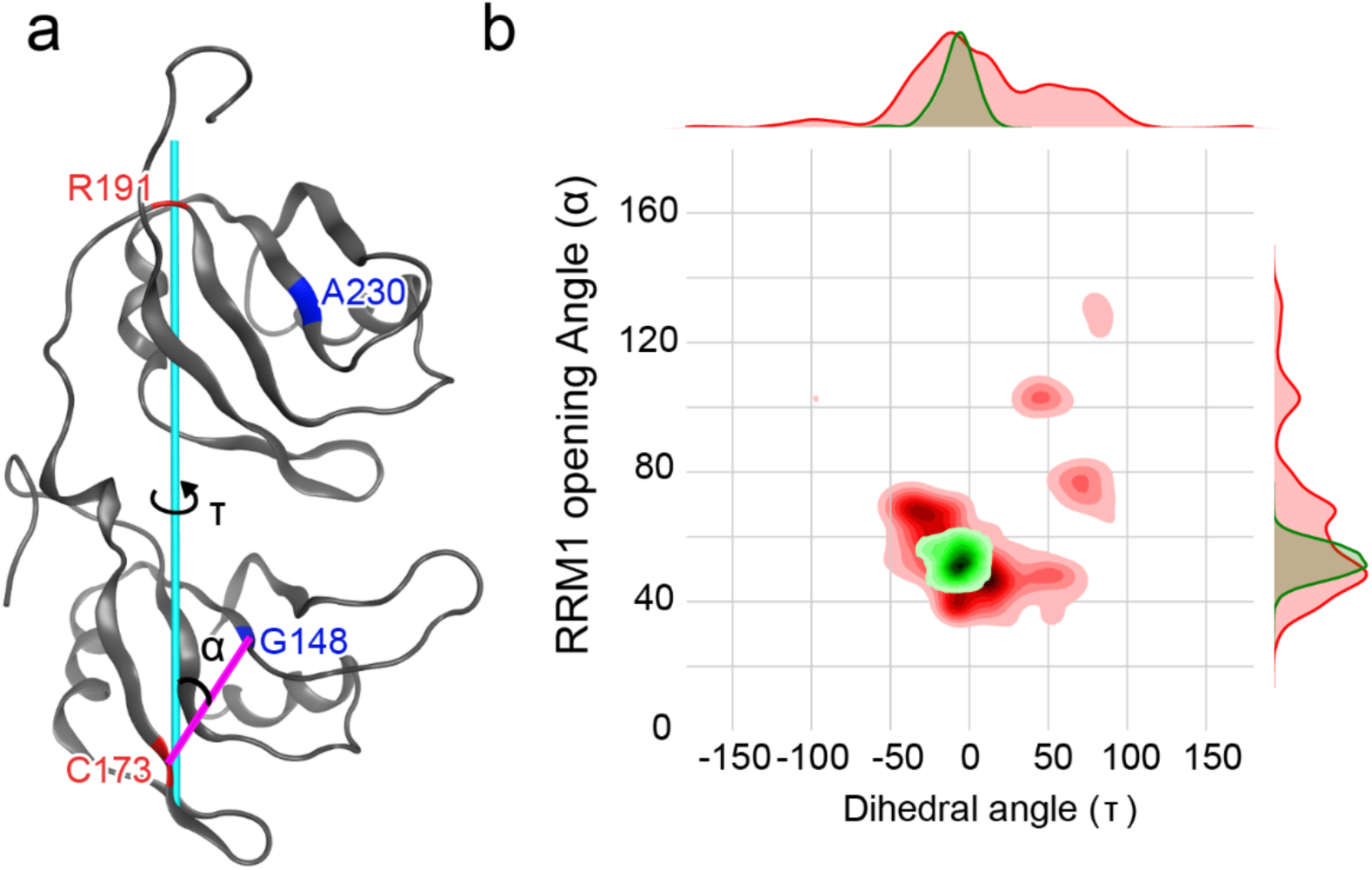
Interdomain angle analysis. (**a**) To measure domain rotation along the axis of the protein, we chose residues G148 and A230 as the center of each domain (blue) and residues C173 and R191 (red) defines the axis of rotation (cyan line). The 4 Cα atoms of these residues defined the dihedral angle (τ) reflecting the relative twisting of these two domains with respect to each other. The RRM1 opening angle was measured using the angle G148 - C173 - R191 (α) centered on Cα atoms of each residue. (**b**) Contour map of the dihedral angle (τ) vs the RRM1 opening angle (α) for RNA-bound (green) and RNA-free (red) simulations. We observe that both angles assume significantly broader distributions in the RNA-free simulations compared with RNA-bound simulations. Furthermore, the RNA-free simulation can be observed in multiple stable conformations.

As a final analysis of dynamic changes between RNA-free and RNA-bound simulations, we analyzed ϕ and ψ angle distributions for each residue over the equilibrated time course (the final 900 of 1000 ns) and analyzed the differences between RNA-free and RNA-bound simulations. We used a 2-dimensional Kolmogorov-Smirnov test for two samples to identify significant changes in ϕ,ψ angle distributions. The only residues showing a significant (p<0.05) difference between RNA-free and RNA-bound simulations were A228 (p=0.006), R227 (p=0.008), E261 (p=0.013), and A260 (p=0.02) (**Fig. 5**). Residues A260 and E261 lie near the C-terminus, while residues R227 and A228 lie at the end of a loop running from I222 to A228. It is worth noting that most changes cluster in the RRM2 domain of TDP-43. The changes in ϕ,ψ angle distribution could be related to changes in the conformation of the (222-228) loop. In the RNA-free system, the loop is able to access a conformation that seems to be restricted by the presence of RNA which brings P223 in closer contact with F231, consistent with our distance analysis.

**Fig. 5.**
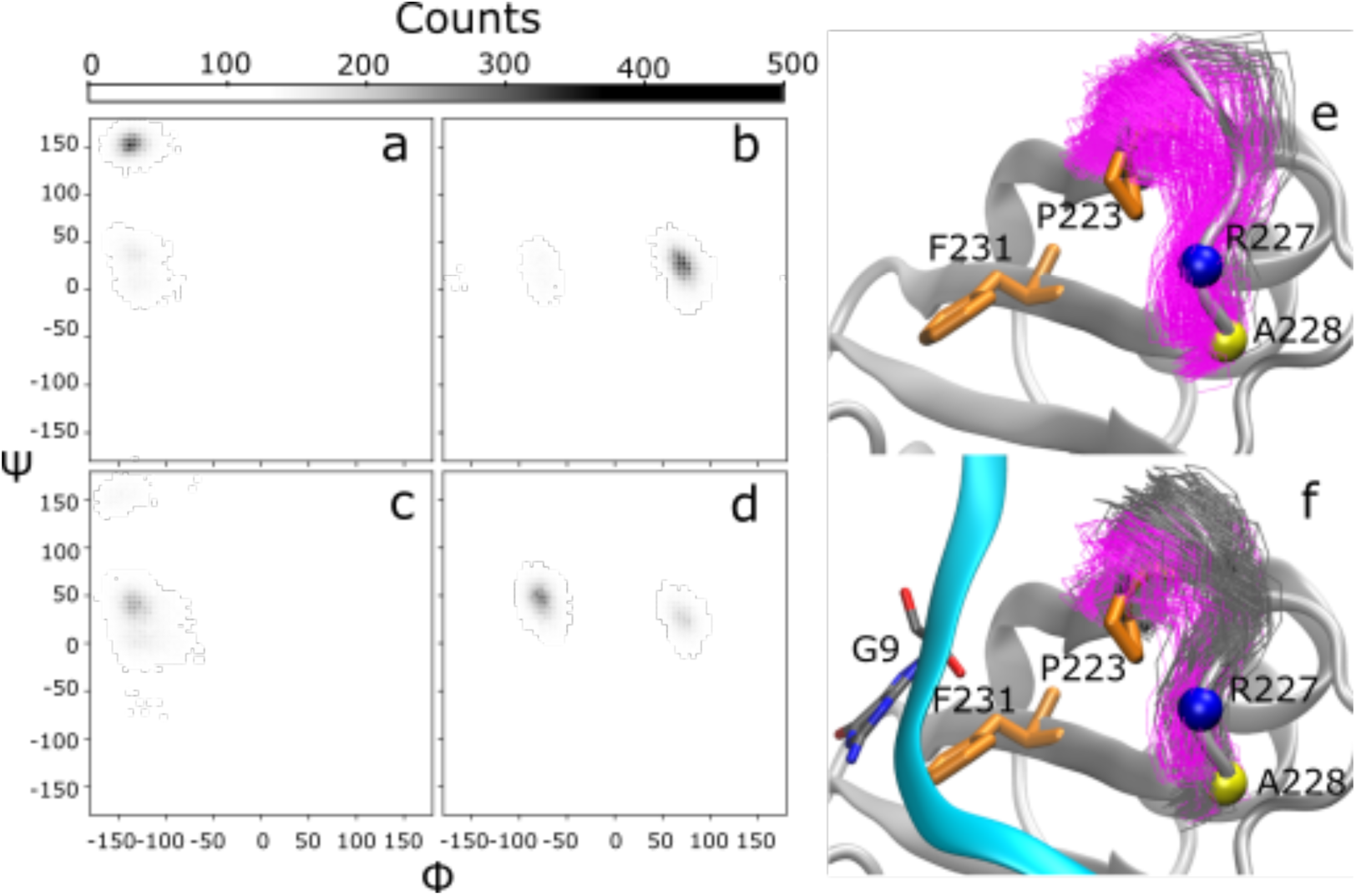
Distributions of phi and psi angles for R227 and A228. Two distinct populations of ψ angles were observed in RNA-free (**a, b**) and RNA-bound (**c, d**) for residues R227 (**a, c**) and A228 (**b, d**) in the simulations of TDP-43. In structure bundles of RNA-free (**e**) and RNA-bound (**f**) TDP-43, loop conformations (residues I222 to A228) are colored grey and magenta corresponding to upper and lower ψ angle clusters of R227, respectively. The larger ψ angle population in RNA-free simulations (lower distribution, magenta) corresponds in a loop conformation shifted towards where RNA would be bound. This movement also corresponds with the shrinking distance between P223 and F231.

### AlphaFold 2 and MD simulation predicted structures and SiteMap Analysis

We compared the top 5 structures predicted by AlphaFold 2. Not surprisingly, the structures of the individual RRM domains were maintained in all models. Those models were aligned to the RNA-bound published structure of TDP-43 (PDB ID: 4bs2^*6*^) as well as to the seven structures derived from the MD simulation (named MD_01 through MD_07) and the corresponding RMSD were analyzed (**Table 1, Fig. S3**). The models AF_2 through AF_5 poorly predicted the 4bs2 structure and any of the MD-derived structures, as the RMSDs ranged from 8.014 to 17.018 Å. However, model AF_1 exhibited the lowest RMSDs (ranging from 2.203 to 5.004 Å) for the RNA-bound RRM domains as well as four different MD derived models, with MD_02 being the closest structure (RMSD = 2.203 Å; **Fig. 6a**). Interestingly, we found that the AF2 predicted TDP-43 RRM domains exhibits structural elements close to either the RNA bound structure or the MD simulated apo structure (**Fig. 6b-f)**. More specifically, the secondary structures α2 of RRM1 (Y155-S162) and β2 of RRM2 (V217-F221) are more similar between AF2 and MD predictions while loops regions (C198-T203 and K136-G146) are more similar between AF2 prediction, and the RNA bound TDP-43 NMR structure. Finally, both MD and AF2 are predicting a longer beta strand in β5 of RRM2 (**Fig. 6b**).

**Table 1.** RMSD for each AlphaFold predicted TDP-43_RRM_ structure compared to 4bs2 and MD derived models.

| **<u>TDP-43 AF1</u>** |  |
| --- | --- |
| 4bs2 | 2.235 (2361 atoms) |
| TDP43_MD_01 | 3.378 (1213 atoms) |
| TDP43_MD_02 | 2.203 (1160 atoms) |
| TDP43_MD_03 | 4.448 (1255 atoms) |
| TDP43_MD_04 | 5.004 (1261 atoms) |
| TDP43_MD_05 | 9.895 (1303 atoms) |
| TDP43_MD_06 | 12.184 (1353 atoms) |
| TDP43_MD_07 | 13.904 (1327 atoms) |

| **<u>TDP-43 AF2</u>** |  |
| --- | --- |
| 4bs2 | 16.476 (2623 atoms) |
| TDP43_MD_01 | 15.283 (1300 atoms) |
| TDP43_MD_02 | 17.018 (1336 atoms) |
| TDP43_MD_03 | 15.412 (1330 atoms) |
| TDP43_MD_04 | 13.606 (1312 atoms) |
| TDP43_MD_05 | 14.047 (1361 atoms) |
| TDP43_MD_06 | 10.655 (1350 atoms) |
| TDP43_MD_07 | 8.014 (1308 atoms) |

| **<u>TDP-43 AF3</u>** |  |
| --- | --- |
| 4bs2 | 14.164 (2648 atoms) |
| TDP43_MD_01 | 12.604 (1321 atoms) |
| TDP43_MD_02 | 14.106 (1325 atoms) |
| TDP43_MD_03 | 12.805 (1356 atoms) |
| TDP43_MD_04 | 10.655 (1348 atoms) |
| TDP43_MD_05 | 12.943 (1318 atoms) |
| TDP43_MD_06 | 9.424 (1352 atoms) |
| TDP43_MD_07 | 8.490 (1309 atoms) |

| **<u>TDP-43 AF4</u>** |  |
| --- | --- |
| 4bs2 | 16.590 (2608 atoms) |
| TDP43_MD_01 | 15.094 (1281 atoms) |
| TDP43_MD_02 | 16.789 (1324 atoms) |
| TDP43_MD_03 | 16.188 (1334 atoms) |
| TDP43_MD_04 | 14.224 (1324 atoms) |
| TDP43_MD_05 | 13.877 (1356 atoms) |
| TDP43_MD_06 | 10.292 (1343 atoms) |
| TDP43_MD_07 | 7.687 (1280 atoms) |
| **<u>TDP-43_AF5</u>** |  |
| 4bs2 | 16.276 (2576 atoms) |
| TDP43_MD_01 | 15.088 (1286 atoms) |
| TDP43_MD_02 | 16.491 (1315 atoms) |
| TDP43_MD_03 | 15.944 (1319 atoms) |
| TDP43_MD_04 | 14.323 (1321 atoms) |
| TDP43_MD_05 | 13.302 (1351 atoms) |
| TDP43_MD_06 | 9.844 (1330 atoms) |
| TDP43_MD_07 | 7.235 (1244 atoms) |

**Fig. 6:**
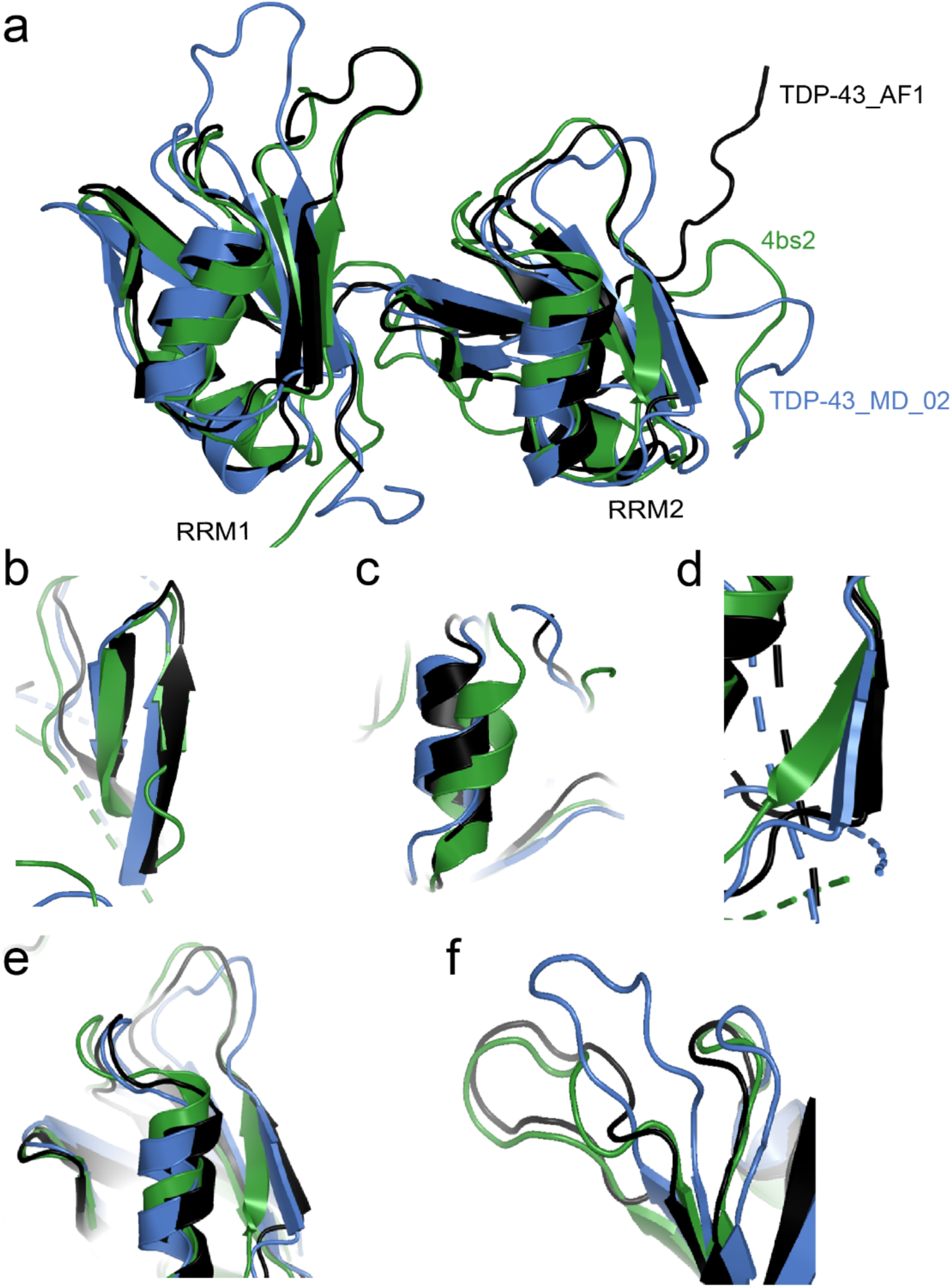
Structural comparison of MD simulated apo TDP-43 with AlphaFold predicted apo TDP-43 and RNA bound known TDP-43 structures. **a**. AlphaFold predicted structure (TDP-43_AF1) was overlayed to published RNA bound (4bs2) and MD derived structure of TDP-43_RRM_ (TDP-43_MD_02). **b**. Zoomed in structure of β4 and 5 in RRM2 (aa. G245-I250). **c**. Zoomed in structure of α2 of RRM1 region (aa Y155-S162). **d**. Zoomed in structure of β2 in RRM2 (aa. V217-F221). **e**. Zoomed in structure of loops in RRM2 (aa. C198-T203). **f**. Zoomed in structure of loop regions in RRM1 (aa. K136-G146).

We then utilized the structures from AF2 and MD and used Sitemap^*16*^ to predict druggable pockets of apo TDP-43 RRM domains to define how dynamics may affect binding sites. Five druggable sites of the average of the RNA bound structure solved by NMR were identified, while only three pockets were found for both MD and AF2 predicted structures (**Fig. S4**). Importantly, a druggable pocket in the central region of the RRM domains that anchors a G-nucleotide (G5 in 4BS2) in the RNA-bound structure was also identified in the MD simulations and AF2-predicted structures. A SiteMap calculation begins with an initial search stage that determines one or more regions on or near the protein surface, called sites, that may be suitable for binding of a ligand to the receptor^*15*^. From this, two important characteristics of pocket interactions can be addressed using SiteScore and DScore. SiteScore determines the probability of ligand-receptor interactions, while DScore determines the druggability of the pocket^*17*^. Our analysis shows that MD structures have poorer SiteScore (0.71 vs 0.779) and DScore (0.58 vs 0.76) compared to AF2 (**Table S1**), indicating AF2 predicted structures may have better pocket formation than MD simulations of apo TDP-43.

## Discussion

Defining targetable pockets of TDP-43 is of therapeutic interest in neurodegenerative diseases. While several targeting approaches have been met with various success, defining which form of TDP-43 will efficiently offer toxicity relief is still being debated. Diversifying the TDP-43 targeting chemical space may bring new opportunities, particularly if small molecules can distinguish RNA-bound and RNA-free form of TDP-43.

Structural determination of the apo RRM domains of TDP-43 has been technically challenging using traditional NMR and X-ray techniques, due to the dynamic nature of the construct. We hence used Molecular Dynamics (MD) simulations starting from RNA bound structure of the tandem domains solved by solution NMR^6^ to sample conformations of the apo TDP-43 structure^*18*^. From the analysis of the simulations, we generally observed an increase in the dynamics upon removal of the RNA as seen in both RMSD and RMSF analysis. Some of the increased dynamics can be explained by additional flexibility allowed to the loops previously contacting the RNA. Notably, the loop spanning residues 138 to 146 showed a particularly large increase in flexibility after the RNA was removed (**Fig. 1**). We also noticed changes in the phi/psi distributions for some residues of the loop_138-146_ were changed, though the change was not as significant as observed for the loop_222-228_. While loop flexibility contributed to the dynamics, most of the additional motion upon removing the RNA came from interdomain motion between RRM1 and RRM2. Contact map analysis reveals that intra-domain contacts are almost unchanged between RNA-free and RNA-bound simulations (**Fig. 2**). However, the RNA-free simulations show a significant decrease in interdomain contacts compared with the RNA-bound simulations. In fact, of the 11 residue pairs with a significant difference between RNA-free and RNA-bound simulations only two (E261-K192 and F231-P223) were not at the interdomain interface and correspond to new contacts observed in the RNA-free versus RNA-bound simulations. The nine remaining residue pairs were all at the interface and showed decreased contact in RNA-free versus RNA-bound simulations. The result is consistent previous observations noting a decrease in inter-domain hydrophobic contacts in much shorter time-scale simulations (100ns)^*19*^. We also note that the result is consistent with experimental evidence of intrinsic low affinity between RRM1 and RRM2 of TDP43 and the requirement of a linker for interaction^*10, 19-22*^.

Looking at the cross-correlation matrix, we detect a significant increase in positive correlation between residue H143 and residues 212 to 217 in the RNA-free vs RNA-bound simulation, as well as significant increases in negative cross-correlation between H143 and residues 129, 152, 153, 194, and 260 in RNA-free vs RNA-bound simulations. Together this is likely due to a more global ‘clam-shell’ movement between the two domains, which appears when the RNA is removed. This ‘clam-shell’ movement is particularly highlighted by our inter-domain angle analysis. In particular, the increased distribution of the α angle in RNA-free simulations highlights this broadening of the angle between RRM1 and RRM2 in RNA-free compared to RNA-bound simulations (**Fig. 4**). The results presented in Figure 4 further serve to highlight the increased sampling of varying orientations between the two domains in the absence of RNA. However, we observe some conformational preference for some states in this plot as indicated by contour peaks where there was greater sampling of a given conformation with respect to neighboring conformations. While the timescales presented in this paper are not likely sufficient to be claim an exhaustive sampling of the accessible conformational space, the use of multiple starting structures and multiple random seeds significantly increases the ability to cover the locally accessible conformational space.

Our interest has been to define druggable sites in TDP-43, either in the RRM or the NTD domain^7,8^. We hypothesized that the RNA-free RRM domains of TDP-43 might exhibit pockets distinct from RNA-bound TDP-43, which could lead to new targeting opportunities. As a first step to drug RNA-free RRM domains, we predicted the apo structure using MD simulations and AlphaFold 2. Sitemap on predicted apo RRM and RNA bound structures revealed fewer pockets in the RNA-free compared to the RNA-bound structures. While this might indicate that small molecules may recognize both RNA-bound and RNA-free, there are more opportunities to specifically target RNA-bound TDP-43. A higher resolution of the RNA-free structure may offer more sites. However, based on our results of the increased dynamics of the apo TDP-43, it may be more challenging to obtain a high resolution of the RNA-free structure.

## Acknowledgements

Research reported in this publication was supported by the Center for Innovation in Brain Science. David Donald Scott is funded by 1T32AG061897-01.

