## Supplemental Data for "Molecular Dynamics simulation of TDP-43 RRM in the presence and absence of RNA"

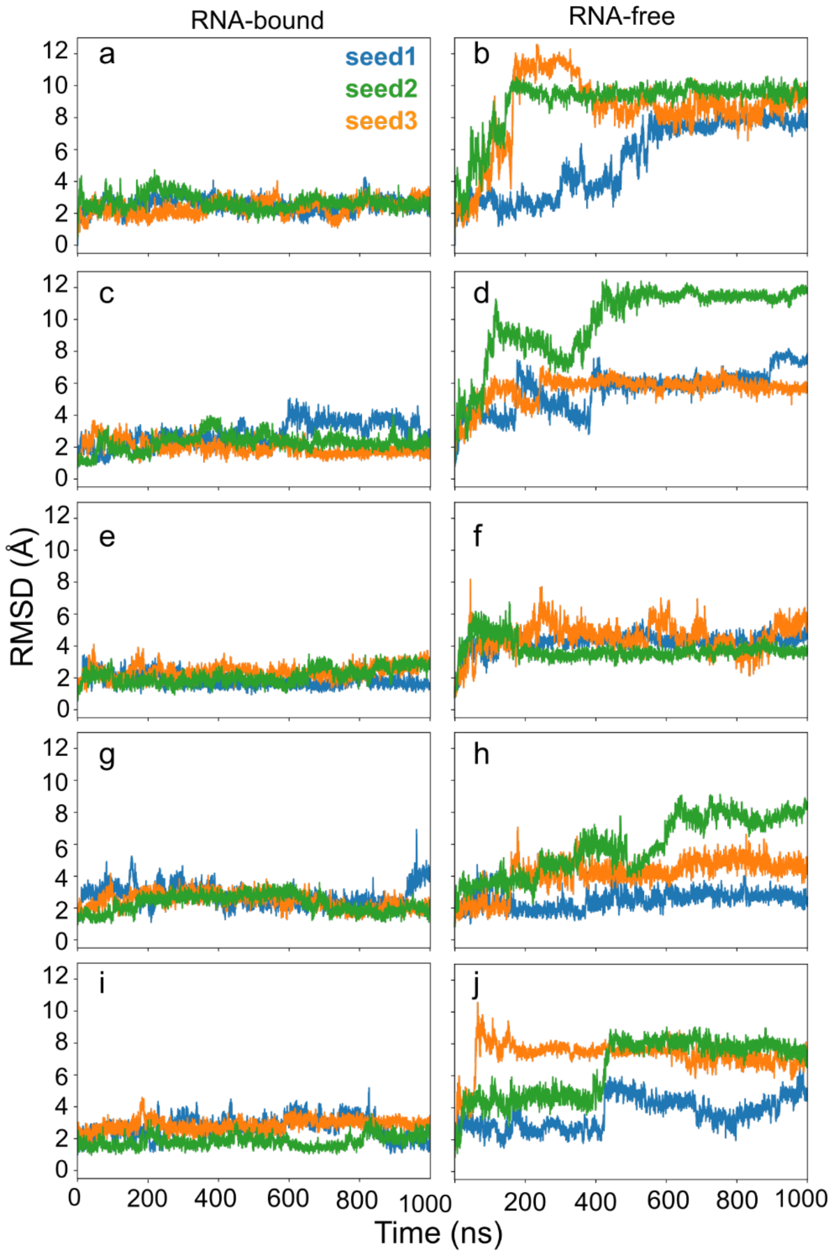

**Figure S1: RMSD of each MD simulation over time.** To evaluate system convergence, RMSD with respect to the reference structure, was computed over the course of the 1-µs simulations. Analysis from simulations starting from each of the 5 sampled NMR structures are segregated by row: (**a**, **b**) conformation 1, (**c**, **d**) conformation 2, (**e**, **f**) conformation 3, (**g**, **h**) conformation 5, and (**i**, **j**) conformation 7. The RNA-bound (a, c, e, g, i) and RNA-free (b, d, f, h, j) simulations are distinguished by columns. The plot color distinguishes the three random seeds (in blue, green and orange). For most (13 out of 15) RNA-bound simulations the trajectories converge within 50 ns of simulation. Seed1 in panels **c** and **e** which show a slight jump later in the trajectory. RNA-free simulations system show much greater deviation from the starting structure and tend to reach equilibrium after 600ns of simulation.

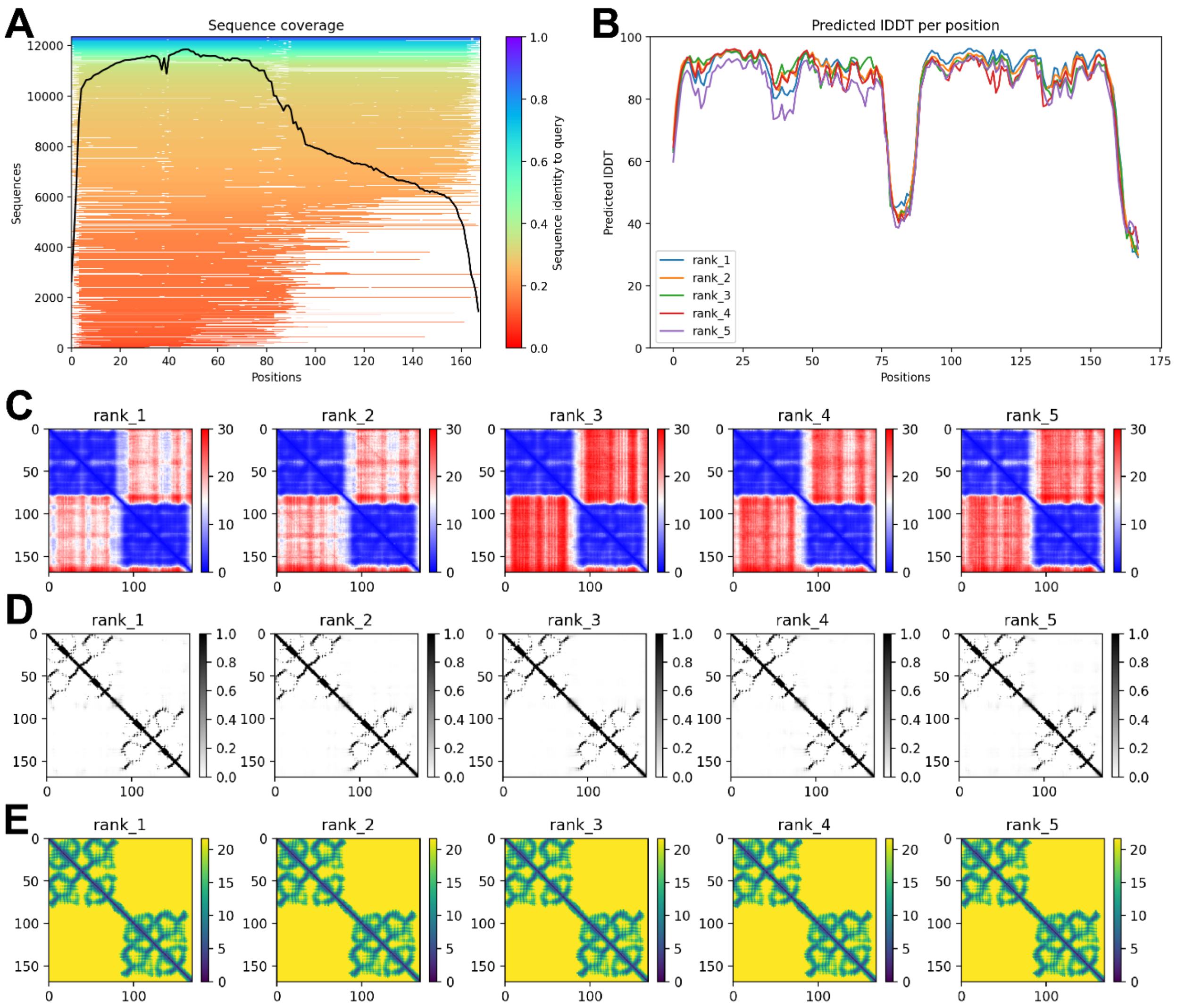

**Figure S2: AlphaFold prediction of TDP-43 RRM domains. A.** Multiple sequence alignment of TDP-43 RRM domains. **B.** Predicted local distance difference test of TDP-43 residues 102-269. **C.** Predicted alignment error of the 5 models obtained. **D.** Predicted contact maps of the 5 models obtained. **E.** Predicted distograms of the 5 models obtained.

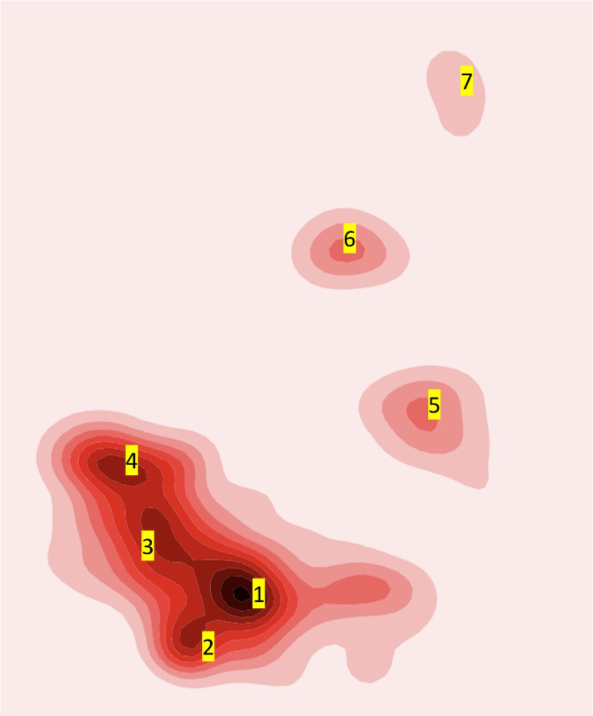

**Figure S3: Contour map of the Md simulated structures showing the seven structures retrieved from the 15 simulations at various different time.**

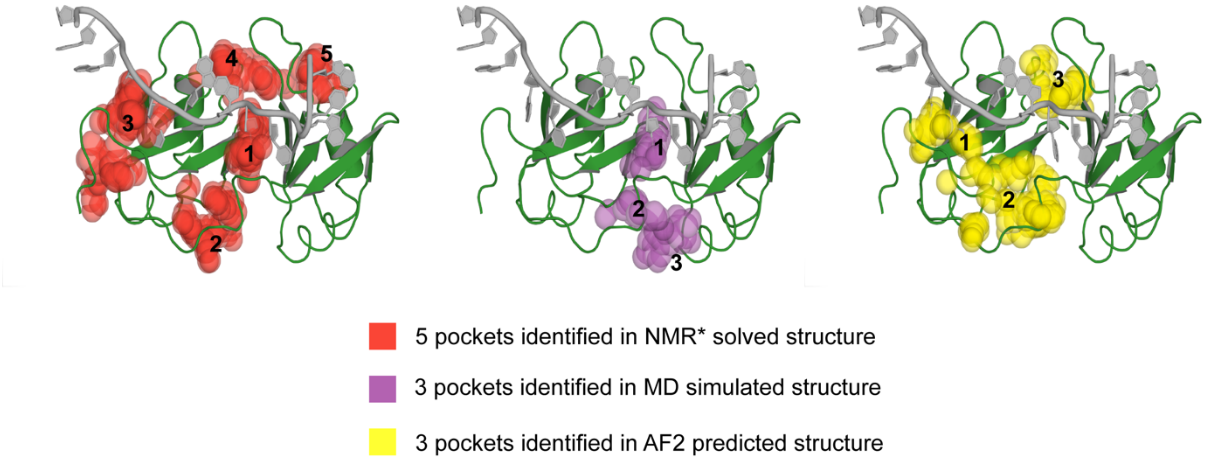

**Figure S4: Sitemap Results of TDP-43 structure models.** Representative image of Sitemap results from all models overlaid onto NMR solved structure PDB 4BS2. Spheres represent contact points within surface pocket density. Data on sites is included in Supplemental Table 1

| Title | SiteScore | Dscore | volume | contact | phobic | philic | balance |
|---|---|---|---|---|---|---|---|
| Ideal scores | ≥ 0.80 | ≥ 1.0 | -- | 1 | 1 | 1 | 1.6 |
| sitemap_RRMs_MD_model2_site_1 | 0.76 | 0.698 | 136.171 | 0.996 | 0.658 | 1.071 | 0.615 |
| sitemap_RRMs_MD_model2_site_2 | 0.656 | 0.575 | 108.045 | 0.855 | 0.223 | 1.079 | 0.206 |
| sitemap_RRMs_MD_model2_site_3 | 0.721 | 0.486 | 136.514 | 1.204 | 0.594 | 1.503 | 0.395 |
| Average MD Sitemap results | 0.712333333 | 0.586333333 | 126.91 | 1.018333333 | 0.491666667 | 1.217666667 | 0.405333333 |
| sitemap_RRMs_AF2_model1_site_1 | 0.87 | 0.862 | 234.269 | 0.803 | 0.197 | 1.08 | 0.182 |
| sitemap_RRMs_AF2_model1_site_2 | 0.746 | 0.742 | 197.911 | 0.642 | 0.054 | 0.895 | 0.061 |
| sitemap_RRMs_AF2_model1_site_3 | 0.721 | 0.678 | 173.901 | 0.789 | 0.166 | 1.063 | 0.156 |
| Average AF2 Sitemap results | 0.779 | 0.760666667 | 202.027 | 0.744666667 | 0.139 | 1.012666667 | 0.133 |
| sitemap_avergae4BS2_site_1 | 0.971 | 0.951 | 219.863 | 0.93 | 0.42 | 1.171 | 0.358 |
| sitemap_avergae4BS2_site_2 | 0.975 | 0.855 | 157.094 | 1.099 | 0.48 | 1.377 | 0.349 |
| sitemap_avergae4BS2_site_3 | 0.872 | 0.905 | 124.166 | 0.904 | 0.836 | 0.77 | 1.085 |
| sitemap_avergae4BS2_site_4 | 0.778 | 0.788 | 122.108 | 0.817 | 0.512 | 0.884 | 0.579 |
| sitemap_avergae4BS2_site_5 | 0.811 | 0.776 | 150.234 | 0.798 | 0.253 | 1.141 | 0.222 |
| Average 4BS2 Sitemap results | 0.859 | 0.831 | 138.4005 | 0.9045 | 0.52025 | 1.043 | 0.55875 |

**Table S1: Sitemap Results for Molecular Dynamics, AlphaFold2, and RNA-bound TDP-43 models.** Sitemap predicted pockets organized by SiteScore, Dscore, volume of pocket and hydrophibicity/hydrophilicity of pocket. Ideal scores are labeled at the top as determined by sitemap calibration. Average results for each model are shown under each subgroup for ease of comparison.
